# *In vitro* determinants of e-cig aerosol condensate bioavailability and toxicity: Influence of cell culture parameters and utility of a normalized dose metric in e-cigarette studies

**DOI:** 10.1101/2021.07.12.452101

**Authors:** Udochukwu C. Obodo, Timothy R. O’Connor

**Author notes:** Corresponding author(s),; (UCO), (TRO).

## Abstract

Electronic cigarettes (e-cigs) have a strong foothold in the marketplace as a product to replace tobacco cigarette usage. Despite many researchers investigating the use of e-cigs and possible health issues, there is still controversy concerning how to evaluate and use e-cig condensates. Therefore, to identify factors that influence *in vitro* e-cig studies, we examined parameters that can impact experimental outcomes. We generated high wattage e-cig aerosol condensate (ECAC) to determine reproducible conditions to evaluate ECAC with respect to cellular survival. Cytotoxicity of ECAC was independent of serum conditions. However, cytotoxicity of ECAC is altered by treatment duration and by physical factors, including cell seeding density and volume of ECAC used. In addition, interactions between ECAC components and cells, as well as the culture vessel surface, diminish the bioavailability of ECAC components *in vitro* and altered the results obtained. Moreover, the cell seeding density changes reactive oxygen species production in response to ECAC exposure. Our data indicated that normalized ECAC doses (ECAC weight per cell) better reflect the toxicity of ECAC than nominal doses (ECAC percentage). These results provide factors for researchers to consider in the design of *in vitro* experiments using ECAC.

## Introduction

Electronic cigarettes (e-cigs) entered the United States marketplace in 2007 [1], and in the years since have rapidly grown in popularity among conventional tobacco cigarette smokers and even non-smokers. As of 2019, an estimated 4.5% of U.S. adults (10.9 million people) [2] were current e-cig users. E-cig aerosols contain significantly diminished levels of carcinogenic compounds relative to tobacco smoke [3–5], thereby mitigating the risk for lung cancer development and/or progression. Nonetheless, there is accumulating evidence that vaping may contribute to other health problems in e-cig users, including non-cancerous respiratory diseases [6–8] and cardiovascular disease [9–11], thus necessitating continued health risk assessment of e-cigs.

Preclinical studies of e-cigs rely heavily, for obvious reasons, on *in vitro* cell culture systems. There is, however, a pervasive lack of consistency in the research methodologies used, ranging from the type of e-cig device used to the methods employed for e-cig aerosol generation and subsequent *in vitro* delivery to cells [12]. These differences can confound the comparisons of results from different studies or laboratories, thereby limiting the combined utility of the generated data. Corrective measures have been proposed or implemented to ameliorate these issues. For example, the National Institute on Drug Abuse (NIDA), in collaboration with NJOY, LLC, developed a standardized research e-cigarette for use in clinical studies to address concerns related to variability in e-cig device design, functionality, and commercial availability [13].

Much less appreciated as potential contributors to variability in *in vitro* e-cig studies are the physical parameters of cell culture, such as cell density and media volume. Cell density and media volume influence cellular response to a number of exogenously delivered substances, including hydrogen peroxide (H_2_O_2_) [14], 1,4-benzoquinone (1,4-BQ) [15], oligomycin A [15], and cigarette smoke extract [16]. In these studies, identical nominal concentrations of the investigative substances, tested under different physical conditions, exhibited varying cytotoxicities that were directly correlated with media volume and inversely correlated with cell density. However, even under different physical conditions, consistent dose-cytotoxicity relationships were obtained for H_2_O_2_, oligomycin A, and 1,4-BQ when media volume and cell number or cellular protein mass (a direct correlate of cell number) were factored in to create an alternative dose metric based on the amount of chemical delivered on a per-cell basis [14, 15]. These findings suggest that for many xenobiotics, nominal concentration may be an inadequate dose metric by failing to account for the contributions from physical parameters in determining the amount of xenobiotic available on a per-cell basis.

Nominal concentrations remain the dose metrics of choice for reporting dose-response relationships in *in vitro* e-cig studies. In only a little over a decade of e-cig research, the reported cellular responses following exposure to e-liquid, aerosol, or any of their components are numerous and varied. These responses include cell death [17–20], reactive oxygen species (ROS) production [21, 22], inflammatory signaling [23–26], mitochondrial dysfunction [19, 27, 28], genotoxicity [18, 20, 29], autophagy inhibition [30], disruption of airway barrier function [31, 32], inhibition of cell differentiation [33, 34], respiratory immune cell dysfunction [35], and dysregulation of DNA repair pathways [36, 37]. Despite this considerable and rapid growth in e-cig literature in the past decade, there are no comprehensive reports to date, and to the best of our knowledge, assessing the potential impact of cell culture parameters on cellular response to e-cig liquid or aerosol exposure. As the e-cig literature continues its meteoric growth, such knowledge is critically needed to determine whether this issue is a potential source of ambiguity that may hamper future cross-study comparisons of *in vitro* toxicology data from e-cig studies. We sought to address this knowledge deficit by exploring the effects of cell culture parameters on cellular response to e-cig aerosol condensate (ECAC). We exposed A549 lung adenocarcinoma cells to ECAC under varying physical conditions and subsequently assessed cell viability using the MTT assay, a widely used method for the assessment of e-liquid and aerosol cytotoxicity. Our goal was to evaluate the utility of nominal concentrations for predicting *in vitro* dose-response relationships following e-liquid or aerosol exposure.

We find that ECAC-induced cytoxicity is dependent on multiple cell culture parameters, including incubation time, cell density, and media volume. Extended incubation of cells with ECAC (up to 24 h) revealed a time-dependent decrease in A549 cell viability. Interestingly, our results also revealed a negative correlation between cell number and ECAC-induced cytotoxicity. Despite being exposed to the same nominal concentrations of ECAC, samples with higher cell counts were better protected from cytotoxicity than those with lower cell counts. In contrast, with other known and controllable variables held constant, ECAC-induced cytoxicity increased with higher volumes of ECAC solution. Furthermore, using a transfer assay developed by Bourgeois *et al.* [16], we show that cell count and media volume are critical *in vitro* determinants of ECAC bioavailability. Finally, we obtained more consistent dose-cytotoxicity relationships with the use of a normalized dose metric (ECAC weight per cell), and briefly discuss the potential utility of this alternative dose metric in advancing *in vitro* e-cig research.

## Materials and Methods

### Cell culture

A549 lung adenocarcinoma cells were kindly provided by the laboratory of Dr. Binghui Shen (Department of Cancer Genetics and Epigenetics, Beckman Research Institute of the City of Hope). Cells were routinely cultured in Dulbecco’s Modified Eagle’s Medium (DMEM) supplemented with 10% fetal bovine serum (FBS), 100 units/ml penicillin, and 100 μg/ml streptomycin, at 37◻C in a humidified incubator containing 5% CO_2_.

### ECAC preparation

DuraSmoke® unflavored e-liquid, containing 36 mg/ml nicotine salt in a base of 50% propylene glycol–50% vegetable glycerin, was purchased from www.Americaneliquidstore.com (Wauwatosa, WI). ECAC was generated inside a fume hood using an Aspire® EVO75 e-cig connected to a SCIREQ inExpose system (SCIREQ, Montreal, Canada) (Fig 1). The e-cig liquid was used with an Aspire 0.5 Ω atomizer and operated at a 70W power setting. The puffing parameters were as follows: 70 ml puff volumes, 3.25 s puff durations, and 60 s inter-puff intervals. Puff timings were controlled by the SCIREQ flexiware software, and the e-cig fire button was manually depressed and released with each puff.

**Fig 1.**
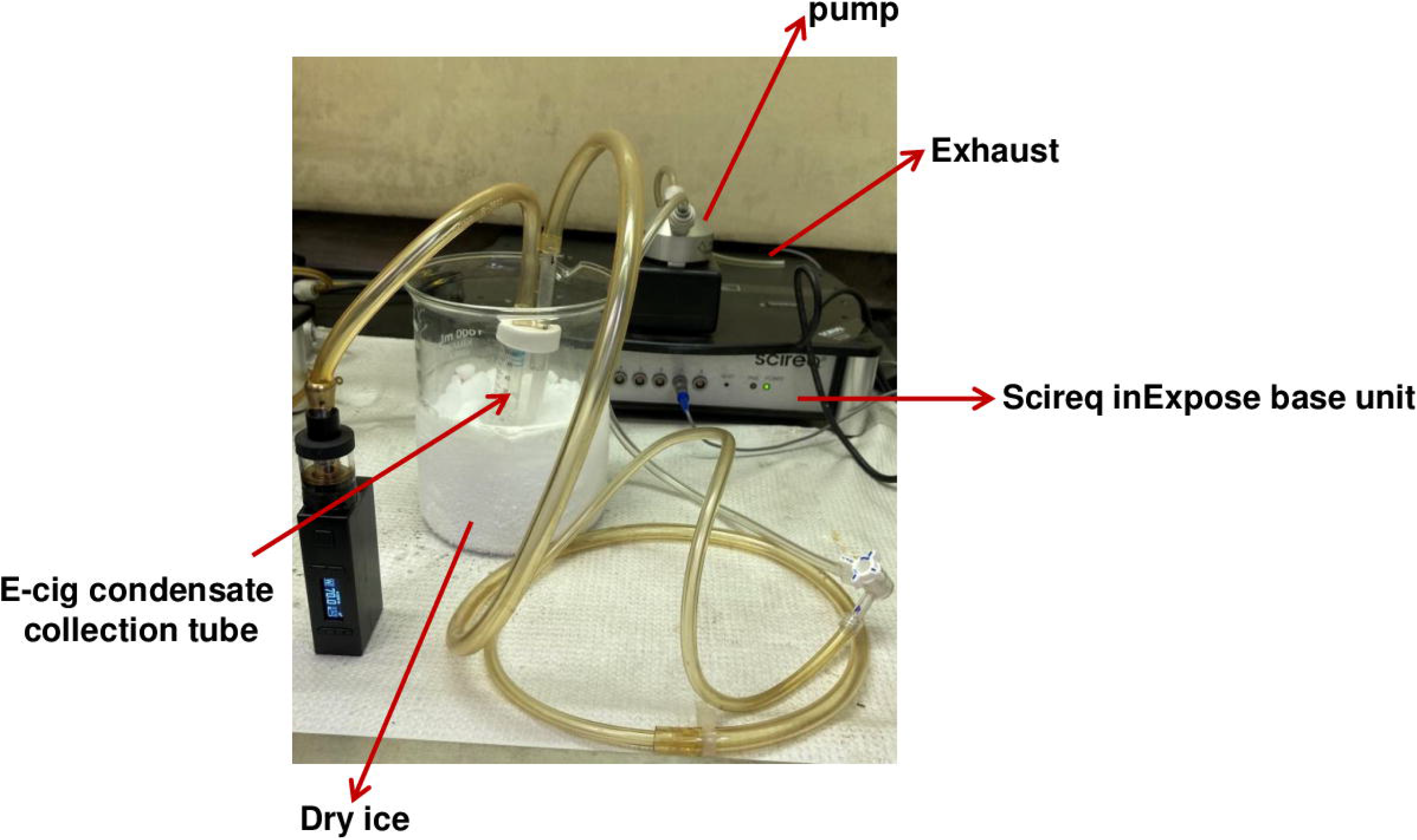
Generation of ECAC. ECAC was generated using the SCIREQ inExpose system. Further details are provided in the Materials and Methods section. Aerosol was collected by condensation in a 50 ml conical tube submerged in dry ice and stored at −80◻C until use.

### ECAC exposures

A549 cells were seeded in 96-well tissue culture plates at the indicated cell densities. Approximately 24 h later, the growth medium was removed, the cell monolayer was washed with calcium/magnesium-free Dulbecco’s phosphate-buffered saline (DPBS), and then incubated for the noted durations with serum-free (unless noted otherwise) DMEM containing ECAC at the indicated doses. Following the desired incubation period, the treatment medium was removed, the cell monolayer was washed with calcium/magnesium-free DPBS and assessments of cell viability and reactive oxygen species (ROS) formation were subsequently performed as described below.

### ECAC transfer assay

A549 cells were seeded at desired cell densities in 96-well plates. Approximately 24 h later, cells in the source wells were pre-incubated with either 150 μl (low volume) or 250 μl (high volume) ECAC at the indicated doses for 2 or 4 h. Following preincubation, 100 μl ECAC was transferred from the low and high volume source wells to destination wells seeded at the onset of the experiment with 15,000 cells. After a 24 h incubation, the MTT cell viability assay was performed as described below. The percent viability of each sample was subtracted from 100% to obtain the percent bioavailability. The formula below was then used to calculate relative bioavailability:

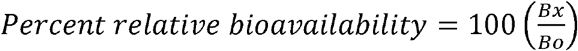

*B*_*x*_ = Bioavailability of each sample

*B*_*o*_ = Average biovailability of the no cell, high volume samples

### ECAC dose normalization experiments

For the ECAC dose normalization experiments, A549 cells were seeded at the desired cell densities in 96-well plates. For each cell density, two extra wells were seeded for the purpose of calculating the number of cells present in the wells at the time of treatment (24 h after seeding) using a hemocytometer-based trypan blue exclusion assay. These pre-treatment cell counts, together with the mass of ECAC (weighed prior to dissolution in media) were used to calculate the normalized ECAC dose in units of ng cell^−1^. The treatments and MTT viability assay were performed as described in the relevant methods section.

### MTT viability assay

MTT solution (Biotium, Hayward, CA) was diluted 1/10 in serum-free antibiotic-supplemented DMEM. One hundred microliters of this dilution were added to the wells of the 96-well plates followed by incubation under normal culture conditions for 3–4 h. After incubation, 100 μl of 10% SDS with 0.01M HCl was added to each well and plates were returned to the incubator for 12 h to solubilize the formazan crystals. Formazan and background optical densities were concurrently measured at 570 nm and 690 nm respectively with a SpectraMax^®^ M3 multi-mode microplate reader (Molecular Devices, San Jose, CA).

### Intracellular ROS assay

ECAC treatments for ROS quantification were performed for 6 h as detailed above with the exception that ECAC was delivered in serum- and phenol red-free DMEM (spf-DMEM). All steps preceding the H_2_O_2_ treatments were as described above for the ECAC exposures. Cells were treated with 10 mM H_2_O_2_ in spf-DMEM for 45 min at 37◻C. At the end of the treatments, cells were washed once with phenol red-free Hank’s Balanced Salt Solution (pf-HBSS) then stained with 0.5 μM CM-H_2_DCFDA dye (Invitrogen, Carlsbad, CA) in pf-HBSS for 40 – 60 min at 37◻C in the dark. After a wash with pf-HBSS to remove excess dye, 100 μl of pf-HBSS was added to each well and fluorescence measurements (485 ±10 nm excitation, 530 ±10 nm emission) were taken on a Cytation3 imaging reader (BioTek, VT). For both ECAC and H_2_O_2_, parallel cultures were treated in the same manner and used for MTT viability assays as described above.

### Statistical analysis

Statistical analysis was conducted in R Studio using two-way linear models, followed by the Holm step-down multiple comparison procedure. P-values < 0.05 were considered statistically significant.

## Results

### High wattage vaping of Durasmoke® e-liquid generates cytotoxic aerosol condensate

The relative cytotoxicities of e-liquids and their corresponding aerosols varies greatly among individual e-liquid formulations, from neither or both eliciting cytoxicity to either the liquid or aerosol, but not the other, displaying cytotoxicity [38]. Therefore, to measure the cytotoxic potential of unvaped Durasmoke® e-liquid versus aerosol, A549 cells were exposed for 24 h to e-liquid or aerosol at doses ranging from 0.25–1% and cell viability was assessed using the MTT assay. Whereas unvaped e-cigarette liquid (ECL) was not toxic to cells under the conditions tested in this assay, e-cig aerosol condensate (ECAC) generated a dose-dependent reduction in cell viability with 34.18% of cells viable at the highest dose (1%) compared to 99.05% viable cells for unvaped e-liquid (Fig 2). Consequently, all subsequent experiments were performed using Durasmoke® ECAC.

**Fig 2.**
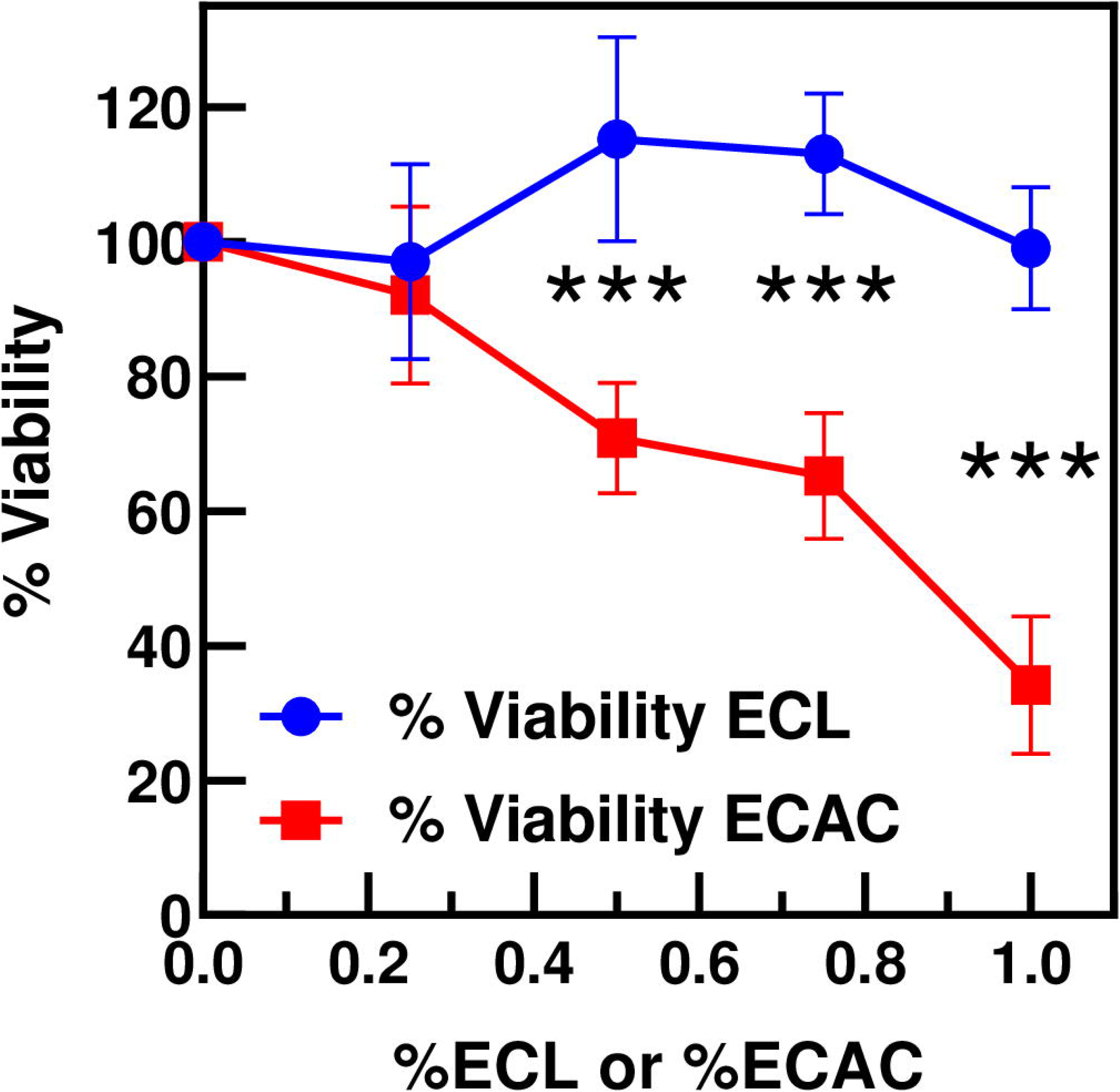
Durasmoke® ECAC, but not ECL, was toxic to A549 cells. A549 cells were seeded at 15,000 cells/well, allowed to grow for 24 h, then treated with ECL or ECAC at the indicated doses for 24 h. The MTT assay was performed immediately after treatment. Cell viabilities are expressed relative to the untreated controls. Data are the averages determined from two independent experiments with triplicate wells per dose level within each experiment. Error bars represent standard deviations (SDs). *** p < 0.001.

### Cytotoxicity of ECAC is independent of serum conditions

The presence of serum in cell culture medium could be a complicating factor in cytotoxicity and genotoxicity assays because of the potential for interactions between serum proteins and the test substances [39–41]. It remains unclear whether serum proteins could influence the cytotoxic potential of ECAC. We addressed this question by exposing A549 cells to ECAC in medium with or without 10% FBS. Two different cell seeding densities were tested (15,000 and 30,000 cells), the first with a low ECAC dose range (0.25–1%; Fig 3A) and the second with a high ECAC dose range (2–3.5%; Fig 3B). The low ECAC dose range resulted in a steady, dose-dependent decline in cell viability (Fig 3A), whereas the high ECAC dose range caused a rapid drop in cell viability with only 25.13% of cells viable at the lowest dose tested (Fig 3B). We hypothesized that any inhibitory effects of serum components on ECAC cytotoxic activity would be more apparent at low to moderately cytotoxic doses (Fig 3A) than at excessively cytotoxic doses (Fig 3B), assuming a mechanism in which serum components bind to and sequester ECAC components from their cellular targets. However, in both circumstances, addition of 10% FBS to the treatment medium had no discernible impact on cell viability. These data suggest that interactions between serum and ECAC constituents are virtually nonexistent or, if present, are either non-consequential or too transient and/or weak to have any tangible effects on ECAC cytotoxic activity.

**Fig 3.**
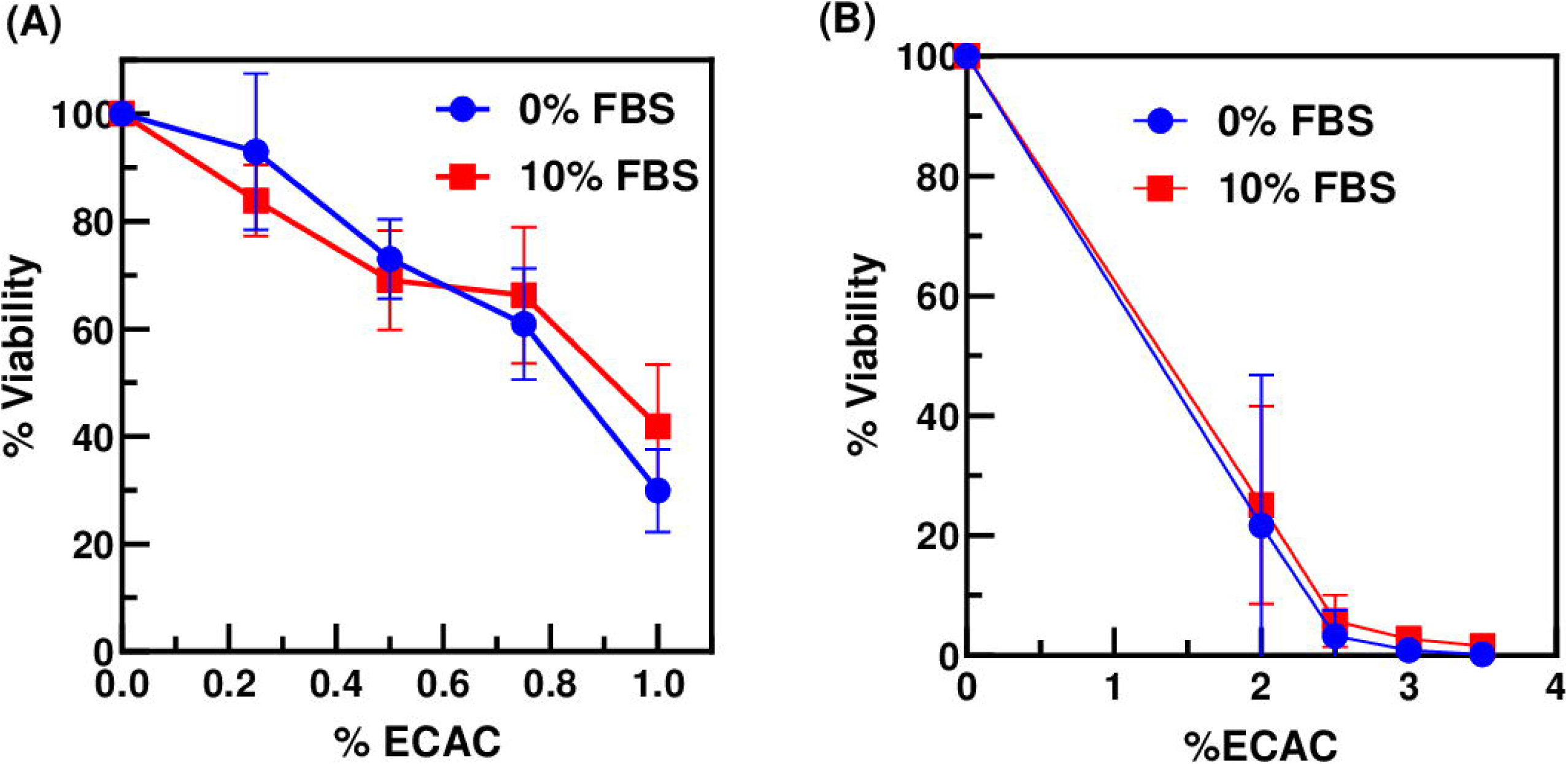
FBS supplementation did not influence ECAC cytotoxic activity. FBS (10%) did not impact Durasmoke® ECAC cytotoxic activity at low to moderately cytotoxic (A) or extremely cytotoxic (B) doses of ECAC. A549 cells, seeded at 15,000 cells/well (A) or 30,000 cells/well (B), were allowed to grow for 24 h, then treated with ECAC at the indicated doses for 24 h. Cell viabilities, measured by MTT assay immediately after treatment, are expressed relative to untreated controls. Data shown are averages ± SDs determined from two independent experiments with triplicate wells per dose level within each experiment.

### Cytotoxicity of ECAC is altered by treatment duration and by physical factors, including cell seeding density and volume of ECAC used

To address the impact of treatment duration on ECAC cytotoxicity, cells (initial seeding density of 15,000 or 30,000 cells/well) were exposed to ECAC (2–3.5%) for 6, 12, or 24 h. Cell viability declined with time, irrespective of the initial cell seeding density (Fig 4A and 4B). However, at the 6 and 12 h time points, ECAC cytotoxicity was blunted when cells were seeded at the higher density of 30,000 cells/well (Fig 4C and 4D), although this protective effect was mostly lost by the 24 h time point (Fig 4E). For example, at the 2% ECAC dose, cell viabilities were 63.93% (15K cells) vs. 87.62% (30K cells) at the 6 h time point, 31.54% (15K cells) vs. 65.52% (30K cells) at 12 h, and 1.90% (15K cells) vs. 13.83% (30K cells) at 24 h (Fig 4C–4E). We observed similar results when cells were seeded at 7,500 or 15,000 cells/well and treated with ECAC at 0.25–1% for the same durations as in Fig. 4. Again, cell viability declined with time for both the low and high cell densities (Fig 4F and 4G). The higher cell density (15,000 cells/well, for this experiment) also conferred increased tolerance to ECAC-induced cytotoxicity at all time points tested (Fig 4H–4J). These data are congruent with previous reports indicating that cell density influences the cytotoxicity of xenobiotics in *in vitro* experiments [14–16].

**Fig 4.**
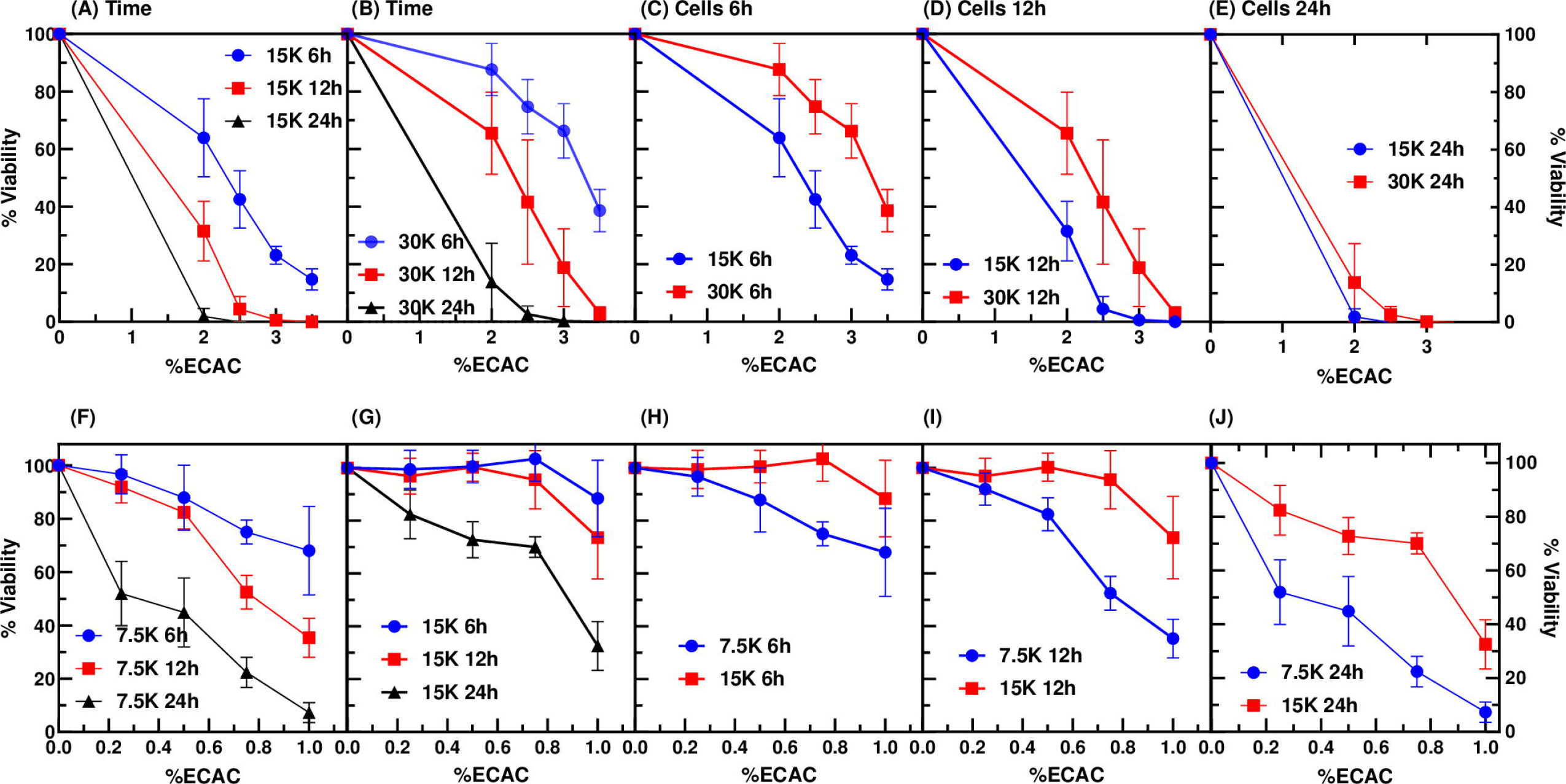
Treatment duration and cell seeding density modulate ECAC-induced cytotoxicity. Durasmoke® ECAC cytotoxicity increased with extended treatment duration and was inversely correlated with cell seeding density. (A–E) A549 cells, seeded at 15,000 or 30,000 cells/well were treated 24 h later with the indicated ECAC doses for 6, 12, or 24 h. (F–J) A549 cells, seeded at 7,500 or 15,000 cells/well, were treated 24 h later with the indicated ECAC doses for 6, 12 or 24 h. Data shown in panels A and B are identical to those shown in panels C–E except that in panels A and B, data are organized so as to isolate the effect of ECAC treatment duration on cell viablity and in panels C–E, data are organized so as to isolate the effect of cell seeding density on cell viability. Data in panels F–J are organized similarly as in panels A–E. MTT assays were performed immediately after treatment to measure cell viabilities, which are expressed relative to untreated controls. Results represent averages ± SDs determined from two independent experiments with triplicate wells per dose level within each experiment.

We also investigated the consequence of varying another physical parameter, the volume of the treatment medium, on ECAC cytotoxicity. Cells (initial seeding density of 15,000 or 30,000 cells/well) were exposed to ECAC (2–3.5%) at three different volumes (50, 100, or 200 μl) for 24 h. Consistent with the data in Fig 4, treatment of the low cell density samples (15,000 cells/well) with 100 μl of the lowest ECAC dose (2%) caused extensive cytotoxicity with only 6.44% of cells viable (Fig 5A). By comparison, 27.98% of cells remained viable following treatment with 50 μl of 2% ECAC and no viable cells were detected after exposure to 200 μl of 2% ECAC (Fig 5A). This inverse relationship between ECAC volume and cell viability was not sustained for all subsequent ECAC doses (2.5, 3, and 3.5%) because maximum cytotoxicity was observed at these doses even for the lowest volume of 50 μl (Fig 5A). In contrast, the high cell density samples (30,000 cells/well) showed a more clearly delineated inverse relationship between ECAC volume and cell viability (Fig 5B). Again, our data are in agreement with previous reports indicating that for a given nominal dose of a xenobiotic, cellular viability generally varies inversely with the volume of the xenobiotic solution [15, 16].

**Fig 5.**
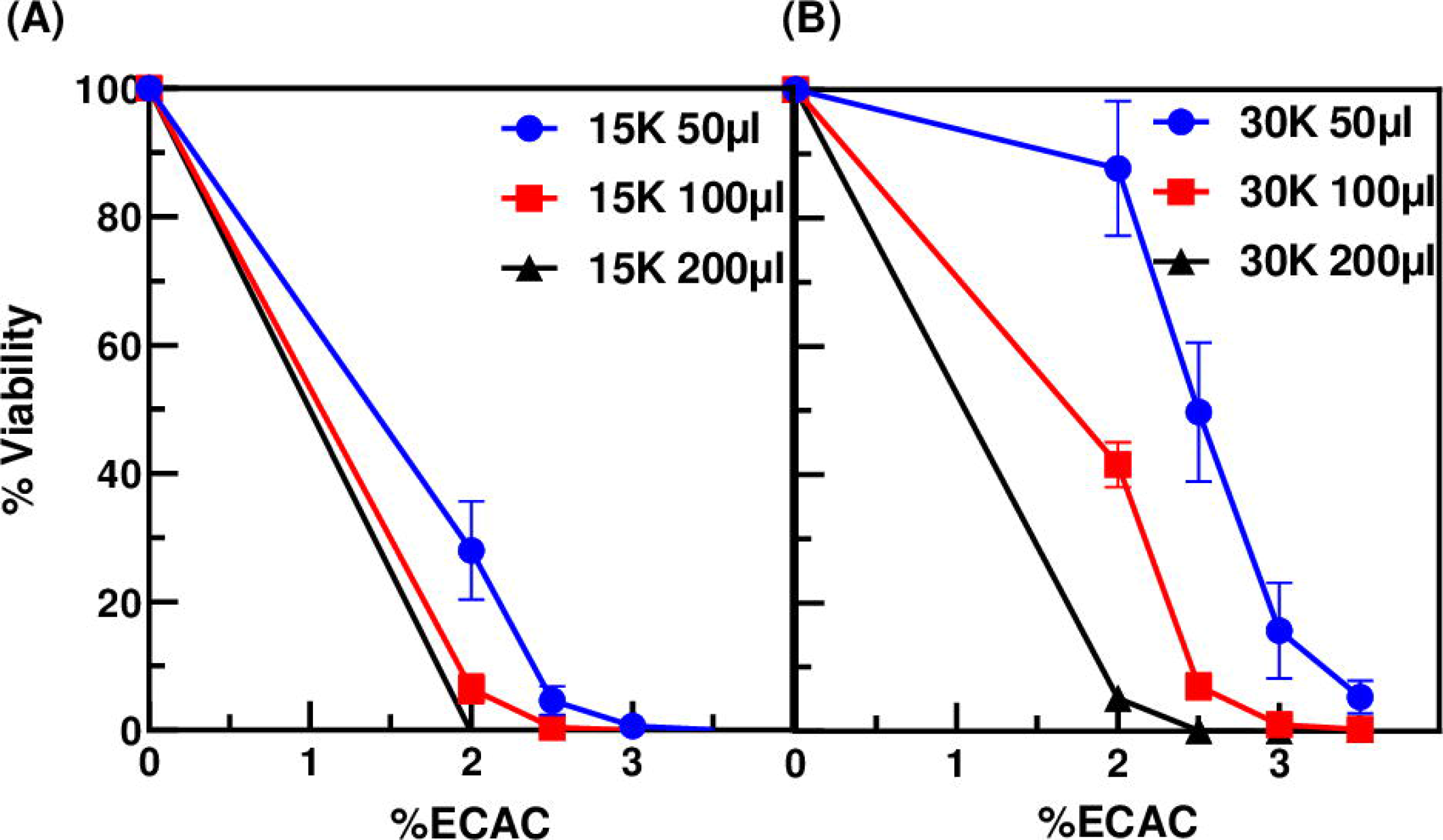
ECAC cytotoxic activity is directly correlated with ECAC solution volume. A549 cells were seeded at 15,000 cells/well (A) or 30,000 cells/well (B), then treated 24 h later with ECAC at the indicated doses and volumes for 24 h. Results of the MTT viability assays are shown with cell viabilities expressed relative to untreated controls. Data are averages ± SDs from two independent experiments with triplicate wells per dose level within each experiment.

### Interactions between ECAC components and cells, as well as the culture vessel surface, diminish the bioavailability of ECAC components *in vitro*

Our results shown in Figs 4 and 5 indicate that the cellular cytotoxic response to ECAC is dependent on the stoichiometric ratio of soluble, bioavailable ECAC components to cells, and suggest that physical factors such as cell density and ECAC solution volume are critical and interrelated determinants of ECAC bioavailability. Bourgeois *et al.* [16] devised a novel assay to ascertain the effects of interactions between soluble cigarette smoke extract (CSE) components and cells on CSE bioavailability. We performed this assay to evaluate the impact of cell seeding density and ECAC solution volume on ECAC bioavailability. A549 cultures at varying cell densities were exposed to a low (150 μl) or high (250 μl) volume of ECAC solution. After the indicated incubation period, 100 μl of ECAC solution was transferred from the source wells to destination wells seeded with 15,000 cells/well and incubated with the new cells for 24 h (Fig 6A). Similar to Bourgeois *et al.* [16], we expressed bioavailability relative to the samples with no cells and with the high volume ECAC solution (0K cells, 250 μl ECAC; Fig 6B).

**Fig 6.**
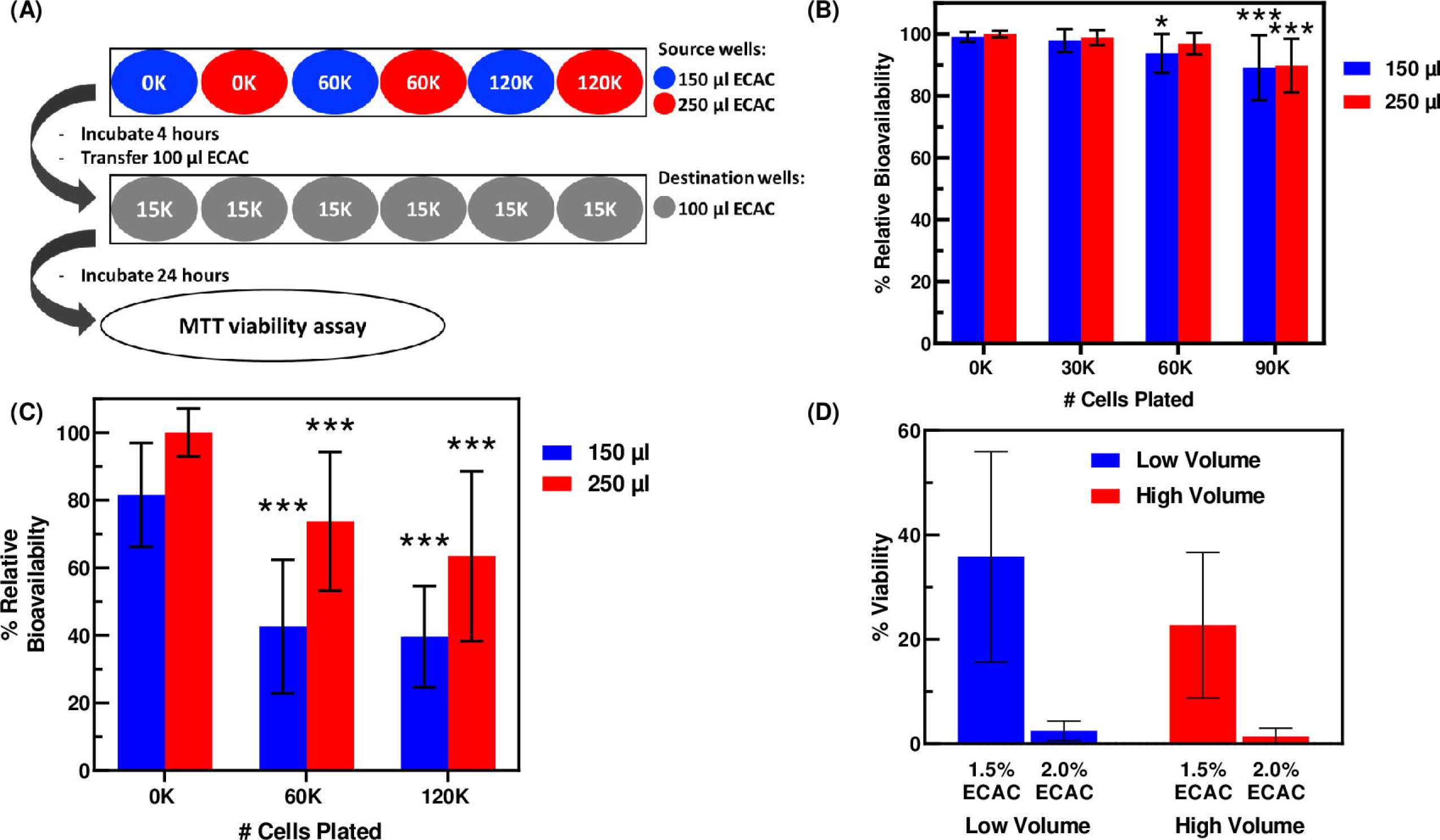
Culture vessel surface, cell seeding density, and ECAC solution volume influence the *in vitro* bioavailability of soluble ECAC components. (A) Schematic of the bioavailability assay. The illustration directly depicts the experiment in panel C but the experiment in panel B was performed in a similar manner. (B) A549 cells were seeded at 0K, 30K, 60K, or 90K cells/well, allowed to grow for 24 h, then treated with 150 μl (low volume) or 250 μl (high volume) of 2% ECAC solution for 2 h. After 2 h, 100 μl of the ECAC solution was transferred from these source wells to destination wells seeded at the onset of the experiment with 15K cells/well and incubated with the cells for 24 h. (C) Experiment was performed in a similar manner as detailed in (B) except that the source wells contained 0K, 60K, or 120K cells/well, ECAC concentration was 1.5%, and the duration of ECAC pre-incubation in the source wells was 4 h. The percent bioavailabilities (calculated as described under “Materials and Methods”) are normalized to the average bioavailability of the 0K high volume samples. Average percent bioavailabilites ± SDs are shown from three independent experiments with 14–16 replicates per data point (panel B) or 24 replicates per data point (panel C). * p = 0.0498, *** p < 0.001 relative to corresponding no-cell control.

In our initial experiment, we used a nominal ECAC dose of 2% and performed the preincubation in the source wells for 2 h (Fig 6B). Under these conditions, we observed a minor though statistically significant effect of cell seeding density on ECAC bioavailability (Fig 6B), presumably because the 2% ECAC dose was extremely cytotoxic in the destination wells (15,000 cells/well; Fig 6D) and the 2 h preincubation in the source wells did not provide ample time for sufficient sequestration of soluble ECAC components prior to transfer to the destination wells. We modified the experiment by using 1.5% ECAC and performing the preincubation for 4 h prior to transferring to the destination plate. In this experiment, we observed a moderate reduction in ECAC bioavailability even in the absence of cells (0K cells, 150 μl ECAC vs. 0K cells, 250 μl ECAC; Fig 6C) suggesting that binding to the culture vessel surface is one route by which ECAC bioavailability could be diminished *in vitro*. For both the low and high volumes of ECAC, incubation with cells resulted in significantly lower bioavailability compared to wells containing no cells. Also of note, at each cell density tested, ECAC bioavailability was better preserved with the high volumes than with the low volumes of ECAC. These data provide strong support for the role of physical factors such as cell density, volume, and culture vessel binding in regulating the *in vitro* bioavailability of soluble ECAC components. Specifically, interactions between ECAC components and cells or the surface of the culture vessel limit the bioavailability of ECAC components, whereas larger ECAC volumes are more resistant than smaller volumes to loss of ECAC components through the aforementioned interactions.

### Cell seeding density also affects the generation of ROS in response to ECAC exposure

Another toxicological endpoint that is frequently measured in preclinical *in vitro* studies of e-cigarettes is ROS production [18, 21, 22, 42]. To investigate whether cell count influences ROS generation, we seeded A549 cells at different densities (15,000–60,000 cells) and exposed them to ECAC (0–3%) or, as a positive control, to H_2_O_2_ (0 or 10 mM). As expected, H_2_O_2_ was toxic to A549 cells (Fig 7A) and also stimulated cellular ROS production (Fig 7B). We validated previous reports of cell count-dependent changes in H_2_O_2_ cytotoxicity [14] (Fig 7A), and report, for the first time to our knowledge, cell count-dependent changes in the stimulation of ROS production by H_2_O_2_ (Fig. 7B). ECAC treatments also induced ROS production, although intracellular ROS levels peaked and then dropped, as cellular viability and metabolism declined, at the highest ECAC doses tested (Fig 7C and 7D). With ECAC treatments, ROS production, like cytotoxicity, was also dependent on cell number. At the seeding density of 15,000 cells per well, maximum ROS induction was observed at the 1% ECAC dose (Fig 7C). In contrast, when cells were seeded at 30,000 cells per well, ROS levels plateaued at the 2% ECAC dose (Fig 7D). In further support of the protective effect afforded by higher cell counts against ECAC-induced ROS generation, peak ROS levels are lower in Fig 7B compared to Fig 7A, despite the 2-fold higher nominal concentration (2% vs 1% ECAC) required to induce maximum ROS production.

**Fig 7.**
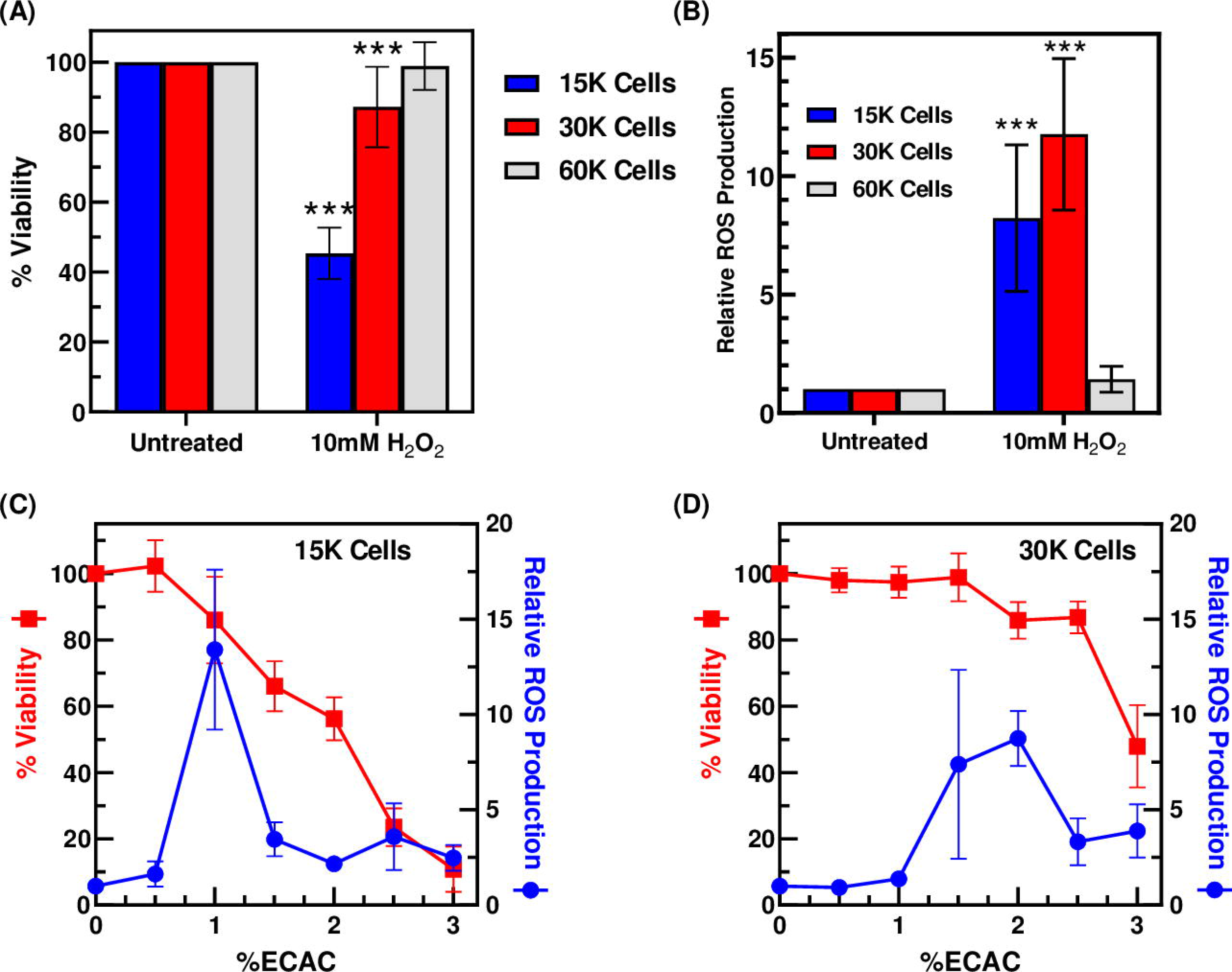
Effect of cell seeding density on H_2_O_2_- and ECAC-induced ROS production in A549 cells. (A) ROS production and (B) percent viabilities following treatment of A549 cells at varying cell densities with H_2_O_2_. (C) ROS production and percent viabilities following treatment of A549 cells (seeded at 15,000 cells/well) with ECAC at the indicated concentrations. (D) The same experiment was performed as in (C) except that cells were seeded at 30,000 cells/well. ROS levels and percent viabilities are normalized to those of the untreated controls. Data are averages ± SDs from three independent experiments, with triplicate wells per data point (panels A and B) or from two independent experiments, with 2–3 replicate wells per data point (panels C and D). *** p < 0.001 relative to corresponding untreated control.

### Normalized ECAC doses (ECAC weight per cell) better reflect the toxicity of ECAC than nominal doses (ECAC percentage)

Our data thus far demonstrate that nominal ECAC doses alone are insufficient determinants of cellular response to ECAC. We tested whether ECAC dose-cytotoxicity relationships may be better represented using normalized doses that reconcile the effects of physical factors like cell count and media volume. This approach has proven successful for other toxicants or xenobiotics. Doskey *et al.* [15] noted that cell count and media volume affect cellular response to oligomycin A and 1,4-BQ, but that these confounding effects were virtually eliminated when drug doses were expressed in units of mol cell^−1^. Similar results were reported by Gülden *et al.* [14] who showed that normalized H_2_O_2_ doses (nmol H_2_O_2_ /mg cell protein) were more reliable predictors of cytotoxic response to H_2_O_2_ than nominal concentrations (molarity). For our purposes, given the multi-constituent nature of e-cig liquids and aerosols [3, 5, 43], we used ECAC weight as the basis for adjustment of the ECAC doses. The ECAC used to prepare the stock treatment solution was weighed prior to dissolution in media. This weight, together with cell counts that were determined prior to treatment, were used to adjust ECAC doses to units of ng cell^−1^. As we demonstrated previously (Fig 4), the data shown in Fig 8A indicate that higher cell counts are more resistant to ECAC-induced cytotoxicity than lower cell counts despite being exposed to the same nominal concentrations of ECAC. This discrepancy was largely resolved when ECAC dose was expressed in units of ng cell^−1^, essentially rendering ECAC cytotoxicity independent of cell count (Fig 8B). Likewise, in experiments where cells were exposed to varying volumes of ECAC, higher ECAC volumes induced greater toxicity (Fig 8C) as previously shown (Figure 5), even though the nominal ECAC dose was constant. In contrast, expressing ECAC dose in units of ng cell^−1^ revealed the expected dose-dependent cytotoxic response (Fig 8D), underscoring the utility of normalized ECAC doses for the accurate representation of ECAC cytotoxic activity.

**Fig 8.**
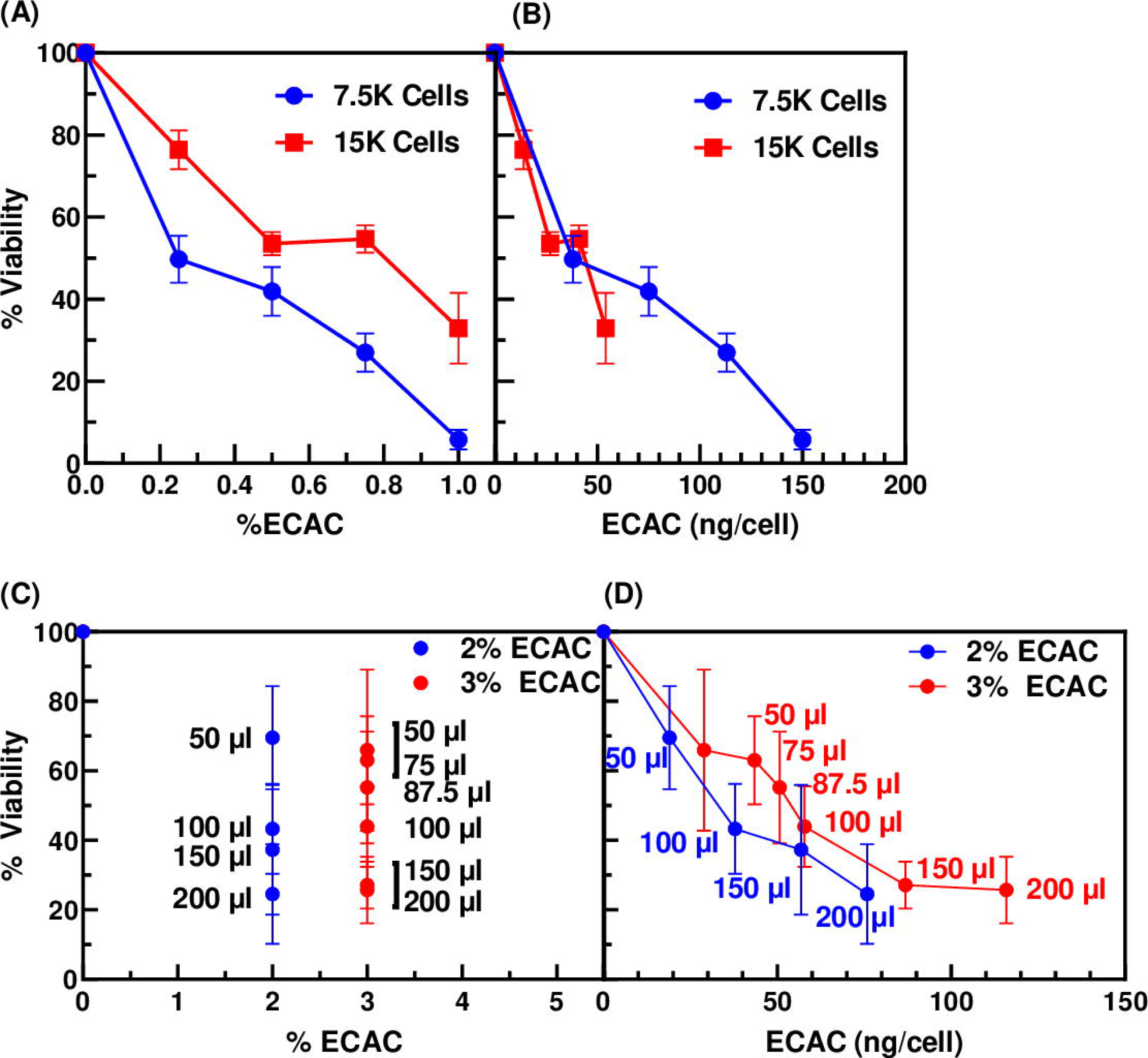
ECAC weight/cell is a more suitable dosing metric for reporting ECAC-induced cytotoxicity than ECAC percentage. (A–B) A549 cells, seeded at 7,500 or 15,000 cells/well, were treated with ECAC for 24 h. Percent viability results in panels A and B are identical, except that ECAC doses are expressed as percentages in panel A and as ECAC weight/cell in panel B. Percent viabilities are averages ± SDs from two independent experiments with triplicate wells per dose level within each experiment. (C–D) A549 cells (30,000/well) were treated with the indicated volumes of 2% or 3% ECAC for 12 h or 6 h respectively. Percent viability results in panels C and D are identical, except that ECAC doses are expressed as percentages in panel C and as ECAC weight/cell in panel D. Percent viabilities are averages ± SDs from two or three independent experiments with 3–6 replicate wells per dose level within each experiment.

## Discussion

*In vitro* assays are particularly advantageous for toxicological testing as they (i) are amenable to high-throughput adaptation, enabling rapid screening of potential toxicants, and (ii) can inform *in vivo* study design and optimization. Dose-response relationships are a fundamental aspect of *in vitro* toxicological testing from which conclusions about the compounds under investigation are drawn. The dosing units used in e-cig studies include percentage [44, 45], total puffs [46], puffs/ml [47, 48], nicotine concentration (molarity or weight/volume concentration) [49, 50], optical density [26], and aerosol particulate matter concentration (weight/volume) [51, 52], all of which are nominal doses. However, nominal dose metrics may be largely ineffective tools for constructing dose-response relationships because they do not incorporate physical parameters, such as cell count and media volume, that are critical determinants of compound bioavailability and cellular response [15, 53]. Gülden et al [14] found this to be the case for hydrogen peroxide, Doskey et al [15] reported similar findings for 1,4-BQ and oligomycin A, and as recently as 2016, Bourgeois et al [16] made similar conclusions about cigarette smoke extract.

Two major aspects in the ongoing contentious debate surrounding the potential health hazards of e-cigs are whether e-cigs are relatively safer than combustible cigarettes, and whether the nicotine and flavorings in e-liquids and aerosols are long-term toxicological hazards. Therefore, studies abound in the e-cig literature aimed at determining the relative toxicological potency of e-cigarette liquids and/or aerosols and cigarette smoke [32, 44, 51, 52, 54], or of different e-liquid formulations or brands [18, 21, 38, 45, 55–57]. However, the use of nominal dose metrics, although seemingly acceptable for these individual studies, may preclude meaningful cross-study comparisons of e-cig toxicology data since experimental conditions vary widely among different studies.

In this study, we evaluated the impact of serum content, incubation time, cell number, and media volume on ECAC-induced toxicity in A549 lung cells. Under our experimental conditions, FBS, at the standard cell culture concentration of 10%, did not interfere with ECAC cytotoxic activity (Fig 3). Serum proteins could compete with cellular targets for binding to the test compounds or could facilitate the cellular uptake of the compounds resulting respectively, in the inhibition or enhancement of cytotoxicity [53]. E-cig aerosols are multi-constituent [3, 5] and their composition depends not only on the initial e-liquid formulation, but also on vaping parameters [58, 59]. Therefore, the nature (or lack thereof) of interactions between e-cig aerosol components and serum proteins will determine how serum influences the cytotoxic potency of individual e-cig aerosol preparations.

In experiments varying the time of exposure to ECAC (up to 24 h), we found that ECAC-induced cytotoxicity increased with extended incubation (Fig 4A, 4B, 4F, and 4G). This phenomenon is hardly unique to ECAC as similar findings have been noted in *in vitro* assays with other chemicals [14, 60, 61]. In e-cig cytotoxicity studies, particularly those involving submerged cell culture models, exposure times are often chosen arbitrarily. Exposure times as short as 1 h [37] and as long as 8 weeks [29] have been reported in the literature. In light of our result that ECAC cytotoxicity is time-dependent, careful consideration should be given to this variable not only during experimental design, but during the review and synthesis of e-cig toxicological data.

We also show, for the first time, that cell number and ECAC solution volume are additional variables that dictate the cytotoxic response to ECAC. The cytotoxic potency of ECAC is inversely correlated with cell number (Fig 4C–4E and 4H–4J), but directly correlated with ECAC solution volume (Fig 5). We note, however, that these relationships are less evident under conditions where ECAC is excessively cytotoxic. For example, when used at the nominal concentrations of 2–3.5%, ECAC cytotoxic potency varies with cell number following 6 and 12 h of treatment (Fig 4C and 4D), but by 24 h, ECAC becomes highly cytotoxic and virtually equitoxic regardless of cell number (Fig 4E). These results suggest that ECAC bioavailability and potency are determined by a complex interplay of exposure time, cell number, and media volume.

We used an assay developed by Bourgeois et al [16] to measure the influence of cell number and media volume on ECAC bioavailability. The two-part assay procedure involves pre-incubating cells in “source” wells with xenobiotic solution for a designated time period, after which xenobiotic solution is removed from the source wells to “destination” wells containing healthy cells for further incubation. Our first iteration of the assay (2 h pre-incubation of 0K, 30K, 60K, or 90K cells with 150 or 250 μl of 2% ECAC solution, followed by transfer of 100 μl ECAC to destination wells) yielded no significant changes to ECAC bioavailability. A potential explanation for this result is that the pre-incubation conditions did not allow for sufficient depletion of the 2% ECAC solution to overcome its exceedingly high initial cytotoxicity (Fig 6D). Our subsequent assay, conducted under less stringent cytotoxic conditions (Fig 6C and 6D), revealed that ECAC bioavailability is reduced by binding to cells and to the culture vessel, although these factors are less impactful for larger ECAC solution volumes than for smaller ones (Fig 6C). It should be noted that this assay does not address the totality of factors that may influence ECAC bioavailability for *in vitro* assays. Evaporation, degradation, and binding to serum components are other avenues by which chemical bioavailability may be reduced *in vitro* [53, 62]. Our bioavailability experiments were conducted under serum-free conditions, although our data in Fig 3 suggest that serum does not affect the bioavailability of our ECAC preparation. Nevertheless, for future studies, such determinations will have to be made for individual e-cig aerosol preparations since e-cig aerosols can differ widely in composition.

Collectively, our results demonstrated that nominal ECAC concentrations are wholly insufficient predictors of cellular response to ECAC exposure. ECAC-induced cytotoxicity and ROS production (Fig 7) were also dependent on physical parameters, including cell number and ECAC solution volume. In lieu of nominal concentrations, Doskey *et al.* [15] propose the widespread use of a normalized dose metric, specified on a per-cell basis, for *in vitro* toxicological assessments. The only information required to compute this metric are the total amount of chemical delivered to the cells, and the cell number at the time of exposure. We showed that specifying ECAC concentration in units of ng cell^−1^ virtually eliminates the dependence of ECAC cytoxicity on physical parameters and is a better approach for reporting ECAC cytoxicity (Fig 8).

Attempts to synthesize the information available from the larger body of e-cig toxicological data are already fraught with complications. The complicating factors include: the sheer number of commercially available e-liquid formulations (over 8,000 unique flavors and counting [63]), differences in e-cig device design and functionality [59, 64], the highly variable vaping protocols used for e-cig aerosol generation [12], and the choice of *in vitro* cell culture platforms available for respiratory research, *i.e.* two-dimensional submerged cultures or three-dimensional air-liquid interface cultures [65]. Calls have been made for the development of standardized research protocols to subvert these issues [12, 66–69]. We propose that such protocols should additionally include standardized dose reporting methods. Nominal and normalized dose metrics, or the pertinent information required to ascertain the latter, including cell number at time of exposure, should be reported. Examples of normalized dose metrics for e-cig studies include weight cell^−1^ (used in this study), puffs cell^−1^, or moles (nicotine) cell^−1^. We anticipate that this approach will improve the combined utility of e-cig toxicological data, allowing for more informative and meaningful conclusions to be drawn from the data, and for the successful translation of *in vitro* data into the design and implementation of animal and clinical studies.

## Conclusion

Physical parameters such as cell number and media volume influence cellular response to e-cig aerosol condensate. Therefore, comparisons between different experimental setups may be challenging in situations where e-cig liquid or aerosol doses are reported only as nominal concentrations. The use of normalized dose metrics, specified on a per-cell basis, can reduce or eliminate this dependency on physical conditions, allowing for meaningful cross-study comparisons of e-cig toxicological data and increasing the amount of information that can be gleaned from the larger body of research on e-cig toxicity.

## Acknowledgments

The authors express their profuse gratitude to Yesenia Thompson for the preparation of the e-cig aerosol condensate used in these studies. We also thank Arnab Chowdhury (Department of Computational and Quantitative Medicine, City of Hope National Medical Center) for performing statistical analysis.

